# The effect of ciprofloxacin and gentamicin on wound healing in *ex vivo* sheep cornea model

**DOI:** 10.1101/2023.11.20.567943

**Authors:** K. Okurowska, D. R. Manrique Suarez, E. Karunakaran

**Affiliations:** Department of Chemical and Biological Engineering, University of Sheffield; National Institute for Health and Care Research, University of Leeds

**Keywords:** *ex vivo* cornea, ocular toxicity, ciprofloxacin toxicity, gentamicin toxicity, delayed corneal wound healing

## Abstract

**Purpose:** Our objective was to assess the efficacy of an *ex vivo* sheep corneal model as an alternative for live animal testing in screening drug cytotoxicity. In pursuit of this goal, we investigated the impact of two commonly used topical antibiotics, ciprofloxacin and gentamicin, on wound healing. Furthermore, we examined different antibiotic dosages and dosing regimens to understand their effects comprehensively.

**Methods:** The epithelium on *ex viv*o sheep corneas was removed with a scalpel, and the area was treated with ciprofloxacin (0.1, 0.3, and 1 mg mL^-1^), gentamicin (0.25, 1, and 3 mg mL^-1^), or phosphate-buffered saline (control). The corneas were exposed to treatments continuously or twice daily for ten minutes. Wound closure was observed by fluorescein retention and histological staining.

**Results:** Untreated corneas healed within 41 hours. Continuous exposure to both ciprofloxacin and gentamicin significantly reduced the corneal healing ability in a time- and concentration-dependent manner. Overall, ciprofloxacin was found to be more toxic than gentamycin. However, this model showed that the corneal epithelium could heal effectively when both antibiotics were administered intermittently.

**Conclusion:** Ciprofloxacin demonstrated greater inhibition of wound healing compared to gentamicin, aligning with *in vivo* studies. The administration of drops several times daily mitigated the toxic effects of antibiotics. The *ex vivo* sheep wound healing model holds promise as an alternative approach to *in vivo* toxicity testing, enabling the swift evaluation of novel antimicrobial treatments and eye drop additives.

## 1. Introduction

The escalating issue of antimicrobial resistance in various pathogens and the subsequent need for novel treatments can be attributed, in part, to the excessive prescription and misuse of antibiotics (Alaali, 2020)Frieri et al., 2017). Despite significant advancements in drug delivery systems, it remains crucial to extensively evaluate the toxicity, absorption, concentration, and effectiveness of both carriers and new compounds before their clinical application (Dubald et al., 2018). Such comprehensive testing is indispensable to ensure the safety and efficacy of these treatments.

Historically, the evaluation of ophthalmic drugs has traditionally relied on the Draize test, involving the direct exposure of live rabbits’ ocular surfaces to drugs (Cormier et al., 1996; Prinsen, 2006; Secchi & Deligianni, 2006). However, this test is lengthy, expensive, and cruel, often resulting in animal suffering and blindness (Lotz et al., 2016; OECD, 2023a). Animals were sacrificed at the end of the experiment. While OECD-validated models exist for measuring opacity and irritants in *ex vivo* bovine (OECD, 2023b) and chicken eyes (OECD, 2023c), the unavailability of eyes from these species of animals from local abattoirs poses a significant limitation. Additionally, the use of chicken eyes for wound healing is hindered by their small size, which would impose severe limitations on the size of wounds that could be studied.

Moreover, the OECD guideline, which assesses acute irritancy and opacity, does not explicitly address wound healing (OECD, 2023a, 2023b). *In vitro* models, on the other hand, are unsuitable for assessing a drug’s effect on wound healing, as they lack the complex physiological processes involved in wound closure, a 3D organ structure, and a limited lifespan (Bu et al., 2019). Therefore, there is a need to find alternatives to live animal testing platforms to test novel ocular drugs (Lieto et al., 2022; Thackaberry et al., 2019).

*Ex vivo* organotypic corneal models have emerged as a promising alternative to *in* vivo models, offering the potential to replace or minimise their use (Urwin et al., 2020). These models involve the removal of the cornea, which is then maintained in an artificial medium that supports crucial tissue functions (Okurowska et al., 2020). Local abattoirs can serve as a valuable source of eyes for these models. The tested drug or chemical can be directly applied to the *ex vivo* cornea, allowing researchers to record any adverse effects of the treatment (Wilson et al., 2015). Studies have demonstrated that *ex vivo* corneal models can accurately assess eye irritation and offer several advantages (Lieto et al., 2022; Prinsen et al., 2017; Prinsen & Koëter, 1993; Spöler et al., 2015; Wilson et al., 2015). However, it is important to note that current *ex vivo* models have limited reproducibility (Lieto et al., 2022), primarily focusing on direct toxicity (Lotz et al., 2016) without addressing delayed wound healing. The wound-healing process can be influenced by various factors such as specific treatments, inflammation, bacterial corneal infections (Caldwell, 2020), and conditions like diabetes (Bu et al., 2019). These factors can complicate recovery, especially after infections or surgeries. Inappropriate therapies can impact the healing process, leading to epithelial defects, discomfort or even visual loss (Ziaei et al., 2018).

This study aimed to investigate the relevance of an *ex vivo* sheep corneal wound-healing model both *in vitro* and *in vivo*. The choice of using sheep corneas was based on their larger surface area compared to chicken or porcine corneas (Greene et al., 2018) and their thicker epithelium (Batista et al., 2016; Lopinto et al., 2017), enabling the creation of significant circular wounds and facilitating the documentation of wound size changes during the healing process. For this study, we selected two commonly used antibiotics, ciprofloxacin and gentamicin, due to their wide application in corneal infections (Mesplié et al., 2009; Watson et al., 2020; Willcox, 2011). It is known that both antibiotics exhibit toxicity at specific concentrations (Alfonso et al., 1990; Tsai et al., 2010) and can potentially delay wound healing (Parmar et al., 2006; Thompson, 2007). Moreover, ciprofloxacin is administered to reduce the risk of infection and promote epithelial closure after refractive or cataract surgery involving an excimer laser (Vijay Zawar & Mahadik, 2014). To evaluate the healing process, fluorescence retention and histological analyses were performed. The proposed model demonstrated its ability to identify ineffective or toxic compounds early in the evaluation process, thus potentially preventing the need for trials on live animals.

## 2. Animals

A total of 32 sheep heads sourced from a local abattoir in the South Yorkshire region were used in this experiment. The sheep were slaughtered for consumption purposes, and their heads were typically discarded as waste. By repurposing these otherwise unused resources, we were able to make efficient use of the available materials for our research.

## 3. Materials and methods

### 3.1. Experimental setup

Fresh sheep heads from a local abattoir in South Yorkshire were used for this experiment. The age of the sheep was between 10 weeks to 6 months. On arrival at the laboratory, the sheep’s eyes were harvested and cultured following a previously developed *ex vivo* porcine cornea infection protocol (Okurowska et al., 2020), with a few alterations described below.

The enucleation procedure was carried out inside a tissue culture cabinet using aseptic techniques and sterile surgical instruments. After removal, the eyeballs were immersed in phosphate-buffered saline (PBS) and decontaminated in a 0.2% povidone-iodine solution for 1 minute.

A 6 mm punch biopsy specimen was gently pressed against the corneal epithelium to mark a perfect circle. The epithelium within the marked area was scraped off using a sterile 15A scalpel blade, creating a wound area with a surface area of 0.280 cm^2^. The cornea was rinsed with 1 mL phosphate-buffered saline (PBS). Two drops of 2% fluorescein sodium salt in distilled water (w/v) were dispensed across the cornea to determine if all epithelium had been removed from the scraped area. The corneas were rinsed twice with 1 mL PBS and photographed using an iPhone 12 Pro Max wide-angle camera secured on a tripod held 12.5 cm above the tissue. Each cornea was lit with a UV torch and photographed next to a ruler for distance calibration. Corneas were separated from the globe as described previously (Okurowska et al., 2020) and left in a medium with antibiotics, while all eyes were processed. Melted agar was used to maintain the shape of the cornea (Okurowska et al., 2020) and secure it to the bottom of a 6-well plate. The corneas were immersed in 3 mL of organ culture medium and incubated as described previously (Okurowska et al., 2020). The composition of the culture medium was: DMEM (phenol-free): F12 (Sigma, Germany) [1:1] supplemented with 5 μg mL^-1^ insulin (SLS, UK) and 10 ng mL^-1^ EGF (SLS, UK), 10% fetal bovine serum (FBS) (Labtech International, UK), 100 U mL^-1^ penicillin, 100 U mL^-1^streptomycin (SLS, UK) and 2.5 μg mL^-1^ amphotericin B (Sigma, Germany). Penicillin and streptomycin were added to eradicate contaminants from the abattoir and typically do not affect corneal migration at this concentration (Nakamura et al., 1993). Amphotericin B was also found to be safe for preserving corneas (Harada et al., 2022). Media were supplemented either with ciprofloxacin hydrochloride, 0.1 (386 μM), 0.3 (778 μM), and 1 (2592 μM) mg mL^-1^ (Sigma, Germany) or gentamicin sulfate, 0.1 (67 μM), 0.25 (168 μM), 1 (672 μM) and 3 (2015 μM) mg mL^-1^ (Sigma, Germany).

Each experiment consisted of two control corneas without antibiotics and two or three treated corneas per condition per week to ensure that the corneas came from a different group of sheep. Some corneas were treated with antibiotics continuously to mimic chronic exposure. In contrast, other corneas were dosed for 10 minutes twice a day, as in the clinical setting (Maggs, 2008), with 0.3 mg mL^-1^ ciprofloxacin hydrochloride and 1 mg mL^-1^ gentamicin. The antibiotic concentrations used were lower than those used in eye drops and were selected based on preliminary testing and toxicity levels from *in vitro* and *in vivo* studies (Alfonso et al., 1990; De Benedetti & Brancaccio, 2010; Sosa et al., 2008; Stevens et al., 1991; Stiebel-Kalish et al., 1998; Tsai et al., 2010). The media were changed twice daily (morning and late afternoon) for four days. The media pH for all treatments was adjusted to 7.4 because the ciprofloxacin solution is acidic. The experiments were repeated three times, aiming for six replicates for each condition. The 6-well plates with prepared corneas were placed on a rocker platform (Figure 40) inside a CO_2_ incubator (Heracell, Thermo Fisher) and incubated with 5% CO_2_ at 37°C.

The rocker was used to mimic blinking to provide nutrients and maintain the corneal surface for longer. The rocking was set up at a 6° angle, 8 rocks per minute (people blink 15-20 times/minute), as described previously (Deshpande et al., 2015). Corneas were stained with fluorescein, rinsed with 1 mL PBS twice and photographed before replacing media as described previously.

### 3.2. Image data analysis

The images of fluorescence-stained sheep corneas were taken in uncompressed format (RAW) at 0, 15, 20, 36 and 41 hours. Adobe Photoshop 2019 (Adobe Inc.) and AutoCAD digital processing software assessed the fluorescein uptake (cm^2^) area using a free-hand lasso tool. The wound area (cm^2^) was photographically evaluated for up to 41 hours in each treatment group. The wound area was measured in pixels and converted into centimetres squared. The images of a ruler placed next to each cornea were used to estimate a pixel-to-centimetres ratio. Some representative images are presented in greyscale (CMYK format) for publication.

### 3.3. Histology

Control corneas were rinsed with 2 mL PBS and fixed overnight in neutral buffered formalin after the wound closure was observed at 41 hours. There were two corneas per condition, and each was prepared on a separate occasion. The tissue was dehydrated in TP1020 (Leica, Germany) and embedded in wax. Afterwards, the cornea was sectioned in the central part at 15 μm with Microtome RM2145 (Leica, Germany), mounted on glass slides and stained with Eosin and Hematoxylin (Merck, UK). Digital images of stained corneas were captured using ProgRes Capture Pro 2.5 (Jenoptik, Germany) and the upright microscope Olympus BX51 (Essex, UK). Scale bars were added on ImageJ 1.53i (USA).

### 3.4. Statistics

Multiple unpaired t-tests were used to assess the statistical analysis of wound closure over time, followed by the Holm-Sidak method using GraphPad Prism version 8.4.1. Statistical significance was set at P < 0.05.

## 4. Results

### 4.1. Control corneas

The study aims to visualise and measure wound closure in the corneal epithelium. To achieve this, we initially confirmed fluorescein staining (Prinsen & Koëter, 1993) in wounded *ex vivo* sheep corneas. Unwounded corneas exhibited a ground glass appearance and a smooth surface on the day of harvesting (Fig. 2A). Despite washing in PBS, corneas with the epithelium removed retained fluorescein, causing them to emit a green fluorescence under UV light. This enabled clear visualisation and measurement of the wounded area (Fig. 2B). The initial surface area of the wound was 0.280 cm^2^.

**Figure 1.**
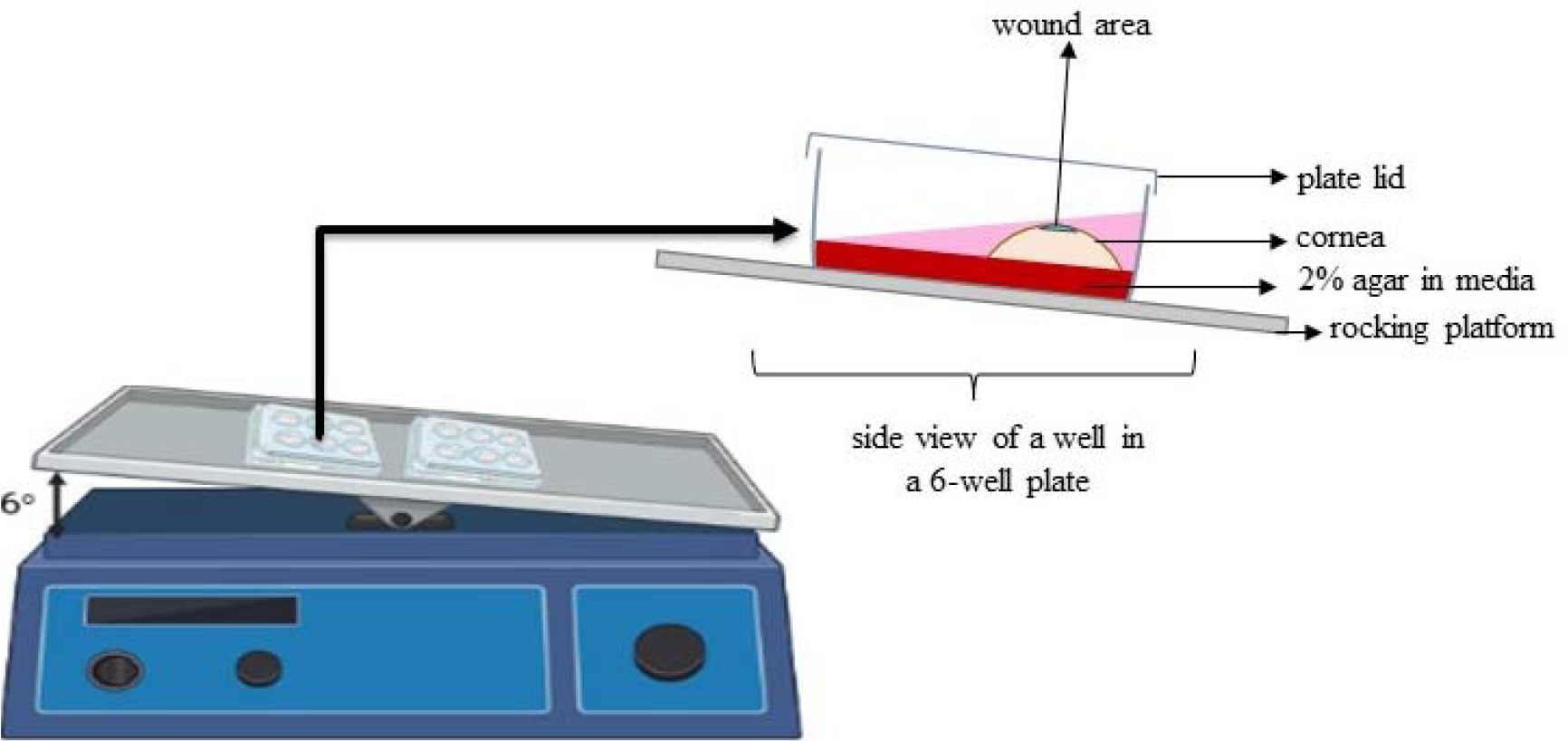
Schematic image of the experimental setup inside a CO_2_ incubator.

**Figure 2.**
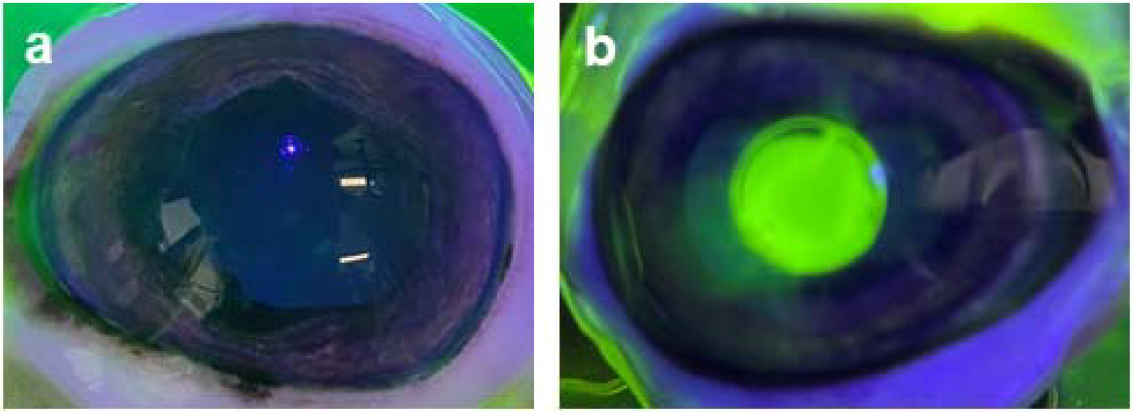
Images comparing intact **(A)** and the sheep cornea with removed epithelium **(B)** followed by staining with 2% fluorescein sodium salt.

### 4.2. Ciprofloxacin treatment

We combined wound measurements from all untreated corneas processed weekly (n = 11). The images showed that eight of the 11 control corneas stopped retaining fluorescein within 41 h (Fig. 3A). The wound size decreased from 0.280 cm^2^ on day 0 to 0.004±0.010 mm^2^ at this time point and was not visible to the naked eye (Fig. 3B).

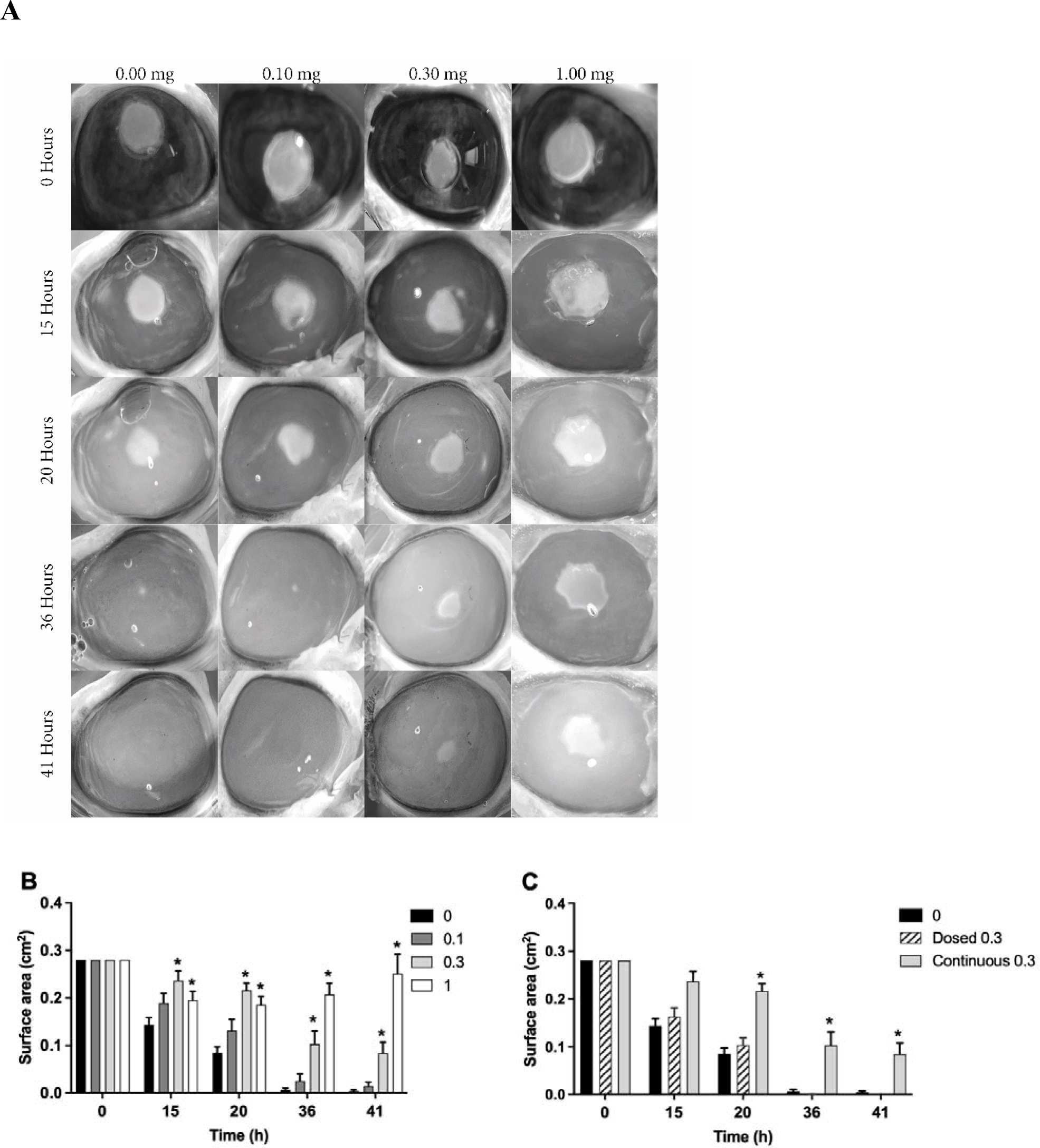

Next, ciprofloxacin was applied continuously at concentrations of 0, 0.1, 0.3, and 1 mg mL^-1^ to wounded untreated and treated *ex vivo* sheep corneas. All corneas treated with ciprofloxacin healed slowly, and a dose-dependent wound healing effect was observed (Fig. 3A and 3 B). Corneas treated with 0.3 and 1 mg mL^-1^ of ciprofloxacin healed significantly slower from 15 h onwards (p < 0.05) (Fig. 3B). At 41 hours, the wound area shrunk with an average diameter of 0.08±0.57 cm^2^ for 0.3 mg mL^-1^ and 0.252±0.091 cm^2^ for 1 mg mL^-1^, respectively (Fig. 3B). This experiment revealed a significant delay (p<0.5) in wound healing at 20, 36, and 41 h after continuous exposure to the highest concentrations of ciprofloxacin. Ciprofloxacin left a white residue in the wound.

Following these results, we applied 0.3 mg mL^-1^ of ciprofloxacin twice a day for ten minutes to see if shorter exposure to this antibiotic would affect wound healing in the same way as a continuous application (Fig. 3C). Corneas healed at the same rate as untreated controls.

These results suggest that continuous exposure to ciprofloxacin concentrations above 0.1 mg mL^-1^ inhibits wound closure in a dose and time-dependent manner. However, this effect was reduced when the drug was applied as an eye drop for a shorter time.

### 4.3. Gentamicin treatment

Next, we continuously applied 0, 0.25, 1, and 3 mg mL^-1^ gentamicin to wounded *ex vivo* sheep corneas and compared them to unwounded control corneas (Fig. 4A and 4 B). All corneas initially healed slowly compared with the controls (Fig. 4B). The wound area decreased significantly from 0.280 cm^2^ (t = 0) to 0.124±0.049 cm^2^ for 1 mg mL^-1^ gentamicin at 20 h (Figure 4B). All corneas healed within 36–41 h. Following earlier results, we increased the concentration of gentamicin to 3 mg mL^-1^ as in eye drops ^28^. Wound healing was inhibited by continuous application of 3 mg mL^-1^ gentamicin at 20 h (0.132±0.048 cm^2^), 36 h (0.051±0.036 cm^2^), and 41 h (0.036±0.035 cm^2^) (Fig. 4B).

**Figure 4.**
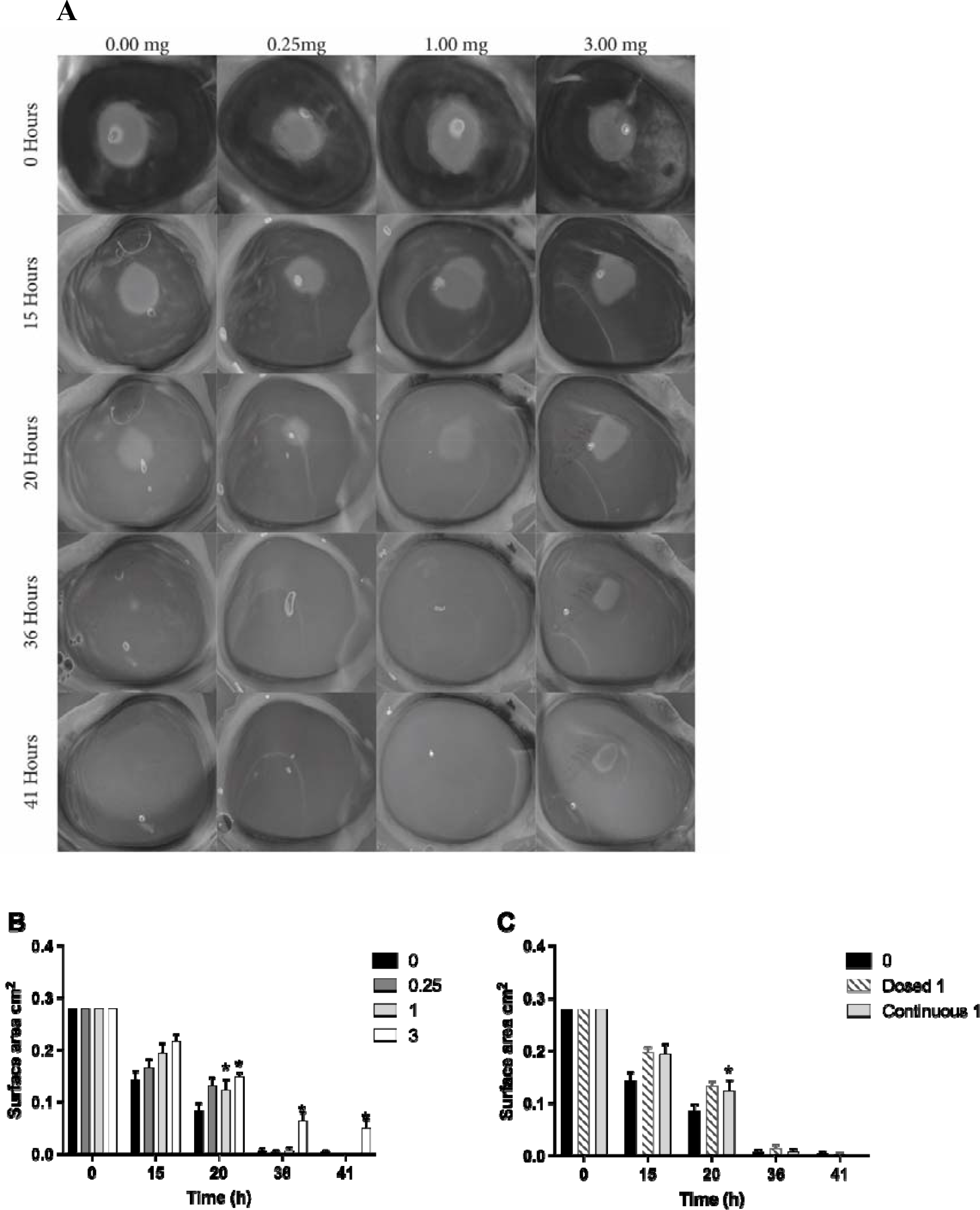
**(A)** Images of fluorescein binding to the wound area in control and treated with gentamicin at 0, 15, 20, 36 and 41 hours. **(B)** Charts show changes in wound diameter (Y axis) over time (X axis) for sheep corneas continuously exposed to gentamicin at concentrations 0 (n = 11), 0.25 (n = 6), 1 (n = 6) and 3 (n = 6) mg mL^-1^ and exposed to 1 (n = 6) mg mL^-1^ of gentamicin twice a day for 10 minutes **(C)**. Mean□ ± □ SEM (*n*□ =□ 6 per group); \**p*□ < □ 0.05 comparing treated vs control, unpaired t-test measures followed by the Holm-Sidak method.

The epithelium in corneas treated with 1 mg mL^-1^ gentamicin twice daily for ten minutes healed slower than that in control corneas (Figure 4C). However, the difference in healing rate was insignificant (p>0.05).

These results suggested that gentamicin is less toxic than ciprofloxacin. Higher doses continuously inhibit wound closure and reduce the harmful effects of antibiotics.

### 4.4. Histology

The tissue sections stained with hematoxylin and eosin revealed a multilayered epithelium in the unwounded sheep corneas. The wounded corneas showed that the epithelium healed at the injury site (Fig. 5B) but appeared thinner than the control epithelium after 41 h (Fig. 5A).

**Figure 5.**
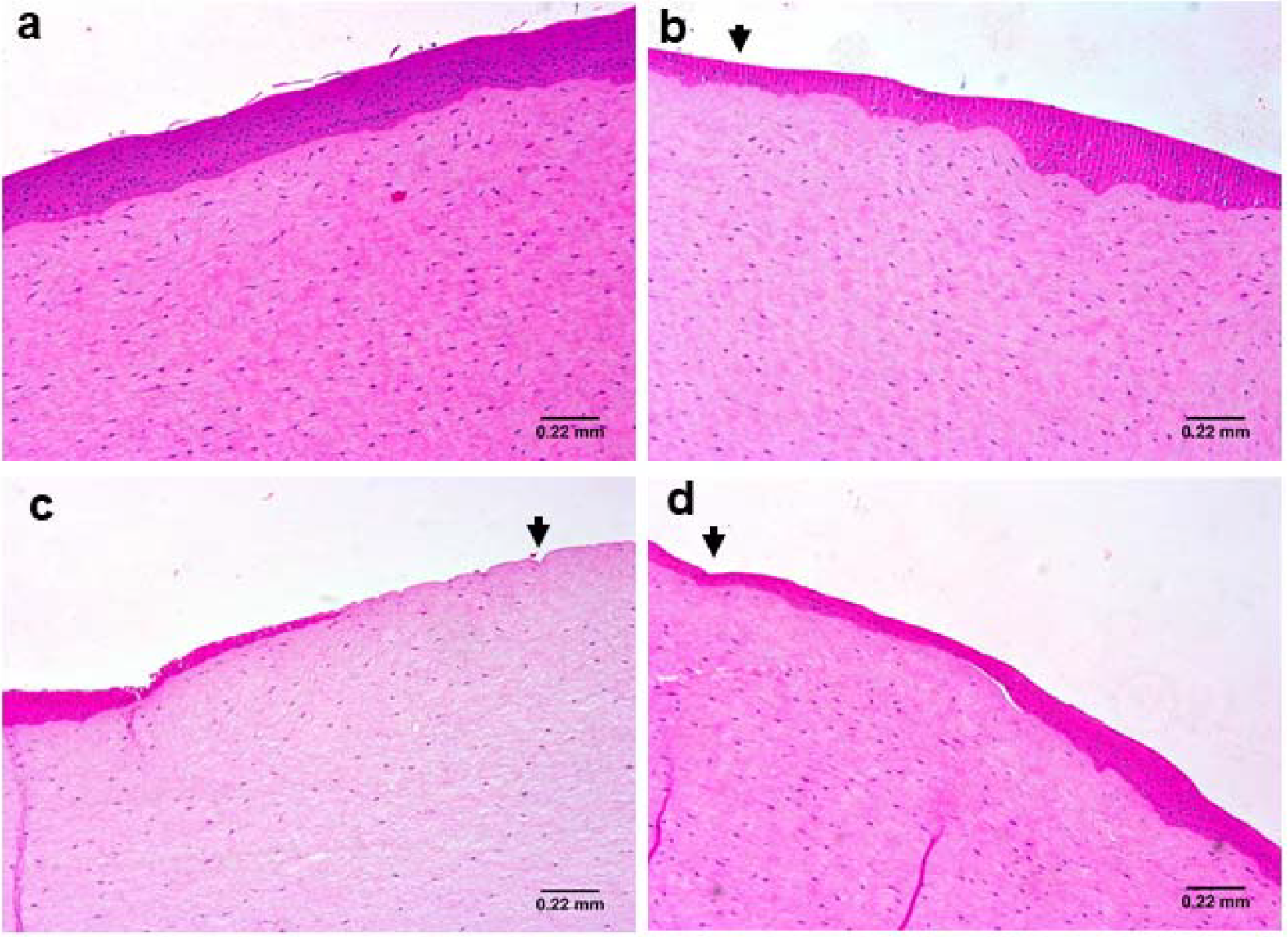
Histology images of corneas after 41 hours of incubation. Unwounded **(A)** and wounded **(B)** *ex vivo* sheep corneas are compared to wounded corneas treated continuously with 1 mg mL^-1^ of ciprofloxacin **(C)** and gentamicin **(D)**. Arrows are pointing at the wound area.

Corneas treated with 1 mg mL^-1^ ciprofloxacin delayed wound healing. The epithelial layer did not regenerate after 41 h, exposing bare stroma. The epithelial cells were rounded, and the nuclei appeared visibly flattened (Fig. 5C). Corneas treated with 1 mg mL^-1^ gentamicin had thinner epithelial layers than the controls at 41 h (Fig. 5D).

## 5. Discussion

In this study, we aimed to investigate the relevance of an *ex vivo* sheep corneal model to study the effect of drugs on the epithelial wound healing process. We used frequently prescribed antibiotics for ocular infections (Watson et al., 2020; Willcox, 2011). Ciprofloxacin is also used post-operation to prevent microbial invasion and enable tissue to heal (Vijay Zawar & Mahadik, 2014). We showed that untreated *ex vivo* sheep corneas healed within 41 hours. As expected, both gentamicin and ciprofloxacin adversely affected the healing rate. This was dose- and concentration-dependent for ciprofloxacin and gentamicin.

In live animals and humans, epithelial cells migrate to the injury site within 24 to 36 hours (Barrientez et al., 2019; Kumari et al., 2018; Wijnholds, 2019) depending on the wound location and size in response to various growth factors and extracellular matrix remodelling (Fini & Stramer, 2005; Ljubimov & Saghizadeh, 2015). The epithelial thickness is fully restored within four weeks (Wijnholds, 2019). Typically, tears supply growth factors to aid the healing process (Klenkler et al., 2007) and this is a limitation of my *ex vivo* cornea model. Studies on *ex vivo* porcine corneas showed that corneal stratified squamous epithelium could regenerate within 72 hours (Schumann et al., 2021) to six days (Castro et al., 2019). In *ex vivo* human corneas, complete reepithelialization was observed after seven days, resulting in five to six layers of the stratified squamous epithelium. The different healing rates observed in *ex vivo* corneal models may be related to several factors. The media composition, size, and depth of the wound and rocking of the cornea during incubation affect the recovery rate (Castro et al., 2019; Deshpande et al., 2015). Our study experiments were terminated once wound closure in untreated corneas was achieved, regardless of the total epithelial thickness.

Since the limbus supplies proliferating cells for cornea regeneration and resides between the sclera’s edge and the cornea (Di Girolamo, 2015; West, 2015), in this study, we preserved some sclera with the limbal region. However, a recent study of *ex vivo* porcine corneas showed that the epithelium without limbus heals at a similar rate (Schumann et al., 2021). Although the limbus region is a source of stem cells, the presence of EGF, insulin and growth factors provided by FBS in media was sufficient to initiate cell replication via mitosis.

Ciprofloxacin was selected for this study because it is used to treat more severe microbial infections of the cornea. The concentration of this drug in eye drops is usually around 3 mg mL^-1^. This concentration showed toxicity and a decrease in cell viability from 30 min of incubation in human corneal epithelial cells *in vitro* (Sosa et al., 2008; Tsai et al., 2010) and severe toxicity after 24 hours in rabbit keratocytes (Seitz et al., 1996). A dose-dependent morphologic change was noted at concentrations as low as 0.003 mg mL^-1^ (Leonardi et al., 2006; Seitz et al., 1996). In our *ex vivo* cornea model, higher than the mentioned concentration *in vitro* (Leonardi et al., 2006; Seitz et al., 1996), 0.1 mg mL^-1^ did not prevent wound closure.

Additionally, our wound healing model allowed for measuring the healing rate over multiple days, providing a more comprehensive understanding than models limited to a few hours. It is worth mentioning that numerous *in vitro* models lack the complexity to replicate the thickness of the corneal barrier (Agarwal & Rupenthal, 2016) and may yield inaccurate predictions regarding drug toxicity levels.

Studies conducted on live rabbits (Moreira et al., 1997) and humans (De Benedetti & Brancaccio, 2010) have demonstrated that re-epithelialization can be delayed depending on factors such as exposure time, presence of wounds, and concentration of ciprofloxacin. Wounding the cornea provides a more relevant representation of clinical scenarios, as antibiotics are typically introduced into damaged corneas after surgery or infection. Ciprofloxacin has been shown to penetrate wounded corneal stratified squamous epithelium more effectively than intact tissue. In live rabbits, the concentration of ciprofloxacin was found higher in the wound area (Fukuda et al., 2004), indicating that drugs can be more toxic in wounded corneas. Injecting 25 μg mL^-1^ ciprofloxacin resulted in corneal swelling and stromal oedema in rabbits (Stevens et al., 1991), with the observed effects being concentration and time-dependent (Stevens et al., 1991). However, it should be noted that antibiotics are typically applied topically to the cornea, not injected. When intact corneas of rabbits were treated with ciprofloxacin (concentration 1.5%) four times a day for three months, no significant ocular irritation or signs of systemic toxicity were observed (Carson et al., 1996). These examples highlight the challenges of comparing results between studies due to variations in experimental procedures employed in *ex vivo* and *in vivo* corneal models. Depending on the specific experimental setup, there is a potential risk of underestimating (Carson et al., 1996) or exaggerating the toxic effects (*in vitro* studies). In a clinical setting, antibiotics are topically used only on corneas with a breached epithelial surface. Therefore, in our study, we ensured the presence of measurable wounds to enhance reproducibility and better replicate clinical conditions.

In clinical practice, administering 3 mg mL^-1^ ciprofloxacin does not typically result in severe side effects for most patients. This antibiotic is usually instilled every 1–2 hours for two days, followed by every 4 hours for 12 days. However, it should be noted that despite this dosing regimen, the use of 3 mg mL^-1^ ciprofloxacin can still lead to discomfort (McDonald et al., 2014) and a slowdown in epithelial regeneration (Patel et al., 2000). The recovery time following ciprofloxacin treatment every six hours can extend up to 120 hours in patients (Patel et al., 2000). It is important to consider that the rate of healing is influenced by factors such as the size of the wound, its depth, and the presence of infection (Schumann et al., 2021).

A small percentage of patients (13-15%) are reported to develop a transient white crystalline corneal precipitate after treatment with ciprofloxacin, which usually resolves within three weeks without treatment (Wilhelmus & Abshire, 2003). However, in some cases, especially in patients with ulcerative keratitis (Parmar et al., 2006; Wilhelmus & Abshire, 2003) or post-surgery (Vijay Zawar & Mahadik, 2014), the precipitate can lead to prolonged epithelial defects (De Benedetti & Brancaccio, 2010; Vijay Zawar & Mahadik, 2014; Wilhelmus & Abshire, 2003). The white precipitation results from the low solubility of ciprofloxacin at the physiologic pH of the tear film (Kanellopoulos et al., 1994). We also observed the precipitation in our study.

Gentamicin demonstrates lower toxicity compared to ciprofloxacin *in vitro* (Tsai et al., 2010). *In vitro* studies have shown a dose- and time-dependent effect of gentamicin (Alfonso et al., 1990; Hendrix et al., 2001). Application of 250 μg mL^-1^ of gentamicin for 48 hours resulted in cytoplasmic granularity in the corneal stratified squamous epithelium (Alfonso et al., 1990). Exposure of human corneal epithelial cells to 1.4 mg mL^-1^ gentamicin showed toxicity within 30 minutes, with decreased cell viability observed after four hours (Tsai et al., 2010). In our *ex vivo* model, concentrations above 1 mg mL^-1^ exhibited mild inhibitory effects on wound healing compared to *in vitro* studies, indicating prolonged exposure may have an impact.

Gentamicin eye drops are commonly prescribed for ocular infections in humans and animals, typically administered 2-12 times daily with concentrations of 3 mg mL^-1^ and higher (Maggs, 2008). However, due to the limited volume of the conjunctival sac and high tear turnover rate (1μL/min), approximately 90% of the eye drop dose is cleared within 2 minutes, necessitating more frequent administration (Maggs, 2008). Studies on rabbits have shown that the corneal stratified squamous epithelium heals with the treatment of 3 mg mL^-1^ of gentamicin. The antibiotic is dosed four times daily during the initial 48 hours and then twice daily (Stiebel-Kalish et al., 1998). A wound diameter of 40 mm^2^ was observed to recover within 96 hours, although the epithelial layer appeared irregular with relatively flat and cuboidal cells (Stiebel-Kalish et al., 1998). Our histology images exhibited a similar effect, with a dose- and concentration-dependent response to gentamicin observed. In our study, sheep corneas treated with gentamicin healed at a comparable rate to untreated corneas, despite using a lower concentration and less frequent dosing than the rabbit study. The differences in dosing and concentration likely account for the more rapid healing observed in our study. Overall, the trend aligned with the findings from the rabbit study.

Data from the wound healing study using the *ex vivo* sheep cornea model holds potential implications for interventions. For instance, cautious monitoring and treatment with ciprofloxacin can be employed to prevent delayed healing and maximise corneal damage caused by the drug in case of infected ulcers. In individuals with ocular surface diseases like dry eye disease, recurrent epithelial erosions, or following extensive corneal surgeries, applying ciprofloxacin and gentamicin may lead to delayed reepithelialisation. Therefore, the dosing frequency and treatment duration should be carefully monitored on an individual basis.

An important limitation of this study is that my *ex vivo* wound healing model employed does not fully replicate the *in vivo* scenario. Factors such as tears, host interactions, and the mechanical removal of drugs by eyelids can influence the drug’s efficacy. Prolonged exposure to the drug can accelerate epithelial damage and exaggerate its impact on wound healing. To address this limitation, antibiotic dosing in this study was designed to mimic clinical conditions, considering these factors.

## 6. Conclusions

The current study highlights the potential value of utilizing the *ex vivo* sheep cornea model to investigate the impact of existing and novel ocular treatments on wound healing. The model offers several advantages, including its cost-effectiveness, ease of setup, and rapid generation of results, which can expedite the screening and development of new drugs. Notably, the model’s ability to measure and observe wound healing progress over time stands as a significant advantage. By allowing modifications in drug dosing regimens and concentrations, this model facilitates the selection of the most effective treatments before progressing to *in vivo* studies.

## Acknowledgements

We express our sincere appreciation to Elliot’s Abattoir in Chesterfield for generously providing the sheep heads used in this study. Additionally, we extend our gratitude to Professor Sheila MacNeil and Professor Annette Taylor for their valuable insights and critical review of this manuscript.

## Author contributions

KO was responsible for the study’s conception and design, acquisition of funding, project management from initiation to completion, data interpretation, and manuscript writing. Both KO and DMS conducted the experiments, with DMS specifically optimising imaging protocols. EK provided valuable feedback on the study design and manuscript.

## Ethical statement

The authors are accountable for all aspects of the work in ensuring the questions related to the accuracy and integrity of any part of the work are appropriately investigated and resolved. Pig eyes were obtained from animals sacrificed for human consumption and not for this study; therefore, ethical approval was not required.

## Funding

The Summer Undergraduate Research Fellowship (SURF) partially sponsored the project.

## Competing Interests

KO, DMS and EK declare no competing interests.

## Data availability

The authors confirm that the data supporting the findings of this study are available within the article.

## Notes

### Competing Interest Statement

The authors have declared no competing interest.

## References

1. Alaali Z, Bin Thani AS. Patterns of antimicrobial resistance observed in the Middle East: Environmental and health care retrospectives. Science of the Total Environment 2020;740.

2. Frieri M, Kumar K, Boutin A. Antibiotic resistance. Journal of Infection and Public Health 2017;10:369–378.

3. Dubald M, Bourgeois S, Andrieu V, Fessi H. Ophthalmic Drug Delivery Systems for Antibiotherapy-A Review. Pharmaceutics 2018;10.

4. Cormier EM, Parker RD, Henson C, et al. Determination of the intra- and interlaboratory reproducibility of the low volume eye test and its statistical relationship to the Draize Eye Test. Regulatory Toxicology and Pharmacology 1996;23:156–161.

5. Prinsen MK. The Draize Eye Test and in vitro alternatives; a left-handed marriage? Toxicology in Vitro 2006;20:78–81.

6. Secchi A, Deligianni V. Ocular toxicology: the Draize eye test. Current Opinion in Allergy and Clinical Immunology 2006;6:367–372.

7. Lotz C, Schmid FF, Rossi A, et al. Alternative Methods for the Replacement of Eye Irritation Testing. Altex-Alternatives to Animal Experimentation 2016;33:55–67.

8. Ubels JL, Clousing DP. In vitro alternatives to the use of animals in ocular toxicology testing. Ocular Surface 2005;3:126–142.

9. Bu YS, Shih KC, Kwok SS, et al. Experimental modeling of cornea wound healing in diabetes: clinical applications and beyond. Bmj Open Diabetes Research & Care 2019;7.

10. Lieto K, Skopek R, Lewicka A, et al. Looking into the Eyes-In Vitro Models for Ocular Research. International Journal of Molecular Sciences 2022;23.

11. Thackaberry EA, Lorget F, Farman C, Bantseev V. The safety evaluation of long-acting ocular delivery systems. Drug Discovery Today 2019;24:1539–1550.

12. Urwin L, Okurowska K, Crowther G, et al. Corneal Infection Models: Tools to Investigate the Role of Biofilms in Bacterial Keratitis. Cells 2020;9.

13. Okurowska K, Roy S, Thokala P, et al. Establishing a Porcine Ex Vivo Cornea Model for Studying Drug Treatments against Bacterial Keratitis. Jove-Journal of Visualized Experiments 2020.

14. Wilson SL, Ahearne M, Hopkinson A. An overview of current techniques for ocular toxicity testing. Toxicology 2015;327:32–46.

15. Spoler F, Kray O, Kray S, Panfil C, Schrage NF. The Ex Vivo Eye Irritation Test as an Alternative Test Method for Serious Eye Damage/Eye Irritation. Atla-Alternatives to Laboratory Animals 2015;43:163–179.

16. Prinsen MK, Koeter H. Justification of the enucleated eye test with eyes of slaughterhouse animals as an alternative to the Draize eye irritation test with rabbits. Food and Chemical Toxicology 1993;31:69–76.

17. Prinsen MK, Hendriksen CFM, Krul CAM, Woutersen RA. The Isolated Chicken Eye test to replace the Draize test in rabbits. Regulatory Toxicology and Pharmacology 2017;85:132–149.

18. Caldwell MD. Bacteria and Antibiotics in Wound Healing. Surgical Clinics of North America 2020;100:757-+.

19. Ziaei M, Greene C, Green CR. Wound healing in the eye: Therapeutic prospects. Advanced Drug Delivery Reviews 2018;126:162–176.

20. Mesplie N, Kerautret J, Leoni S, Dubois V, Colin J. Severe bacterial keratitis and activity of fluoroquinolones. Journal Francais D Ophtalmologie 2009;32:273–276.

21. Willcox MDP. Review of resistance of ocular isolates of Pseudomonas aeruginosa and staphylococci from keratitis to ciprofloxacin, gentamicin and cephalosporins. Clinical and Experimental Optometry 2011;94:161–168.

22. Watson SL, Gatus BJ, Cabrera-Aguas M, et al. Bacterial Ocular Surveillance System (BOSS) Sydney, Australia 2017-2018. Communicable Diseases Intelligence 2020;44.

23. Alfonso EC, Albert DM, Kenyon KR, Robinson NL, Hanninen L, D’Amico DJ. In vitro toxicity of gentamicin to corneal epithelial cells. Cornea 1990;9:55–61.

24. Tsai TH, Chen WL, Hu FR. Comparison of fluoroquinolones: cytotoxicity on human corneal epithelial cells. Eye 2010;24:909–917.

25. Thompson AM. Ocular toxicity of fluoroquinolones. Clinical and Experimental Ophthalmology 2007;35:566–577.

26. Parmar P, Salman A, Kalavathy CM, et al. Comparison of topical gatifloxacin 0.3% and ciprofloxacin 0.3% for the treatment of bacterial keratitis. American Journal of Ophthalmology 2006;141:282–286.

27. Zawar SV, Mahadik S. Corneal deposit after topical ciprofloxacin as postoperative medication after cataract surgery. Canadian Journal of Ophthalmology-Journal Canadien D Ophtalmologie 2014;49:392–394.

28. Gelatt KN. Ocular pharmacology and therapeutics. Essentials of Veterinary Ophthalmology, 3rd Edition 2014;66–99.

29. Stevens SX, Fouraker BD, Jensen HG. Intraocular safety of ciprofloxacin. Archives of Ophthalmology 1991;109:1737–1743.

30. De Benedetti G, Brancaccio A. Corneal Deposit of Ciprofloxacin after Laser Assisted Subepithelial Keratomileusis Procedure: A Case Report. Journal of Ophthalmology 2010;2010.

31. Sosa AB, Epstein SP, Asbell PA. Evaluation of toxicity of commercial ophthalmic fluoroquinolone antibiotics as assessed on immortalized corneal and conjunctival epithelial cells. Cornea 2008;27:930–934.

32. Alfonso EC, Albert DM, Kenyon KR, Robinson NL, Hanninen L D.J. DA. In vitro toxicity of gentamicin to corneal epithelial cells. Cornea 1990;55–61.

33. Stiebel-Kalish H, Gaton DD, Weinberger D, Loya N, Schwartz-Ventik M, Solomon A. A comparison of the effect of hyaluronic acid versus gentamicin on corneal epithelial healing. Eye 1998;12:829–833.

34. Deshpande P, Ortega I, Sefat F, et al. Rocking Media Over Ex Vivo Corneas Improves This Model and Allows the Study of the Effect of Proinflammatory Cytokines on Wound Healing. Investigative Ophthalmology & Visual Science 2015;56:1553–1561.

35. Wijnholds J. “Basal Cell Migration’’ in Regeneration of the Corneal Wound-Bed. Stem Cell Reports 2019;12:3–5.

36. Barrientez B, Nicholas SE, Whelchel A, Sharif R, Hjortdal J, Karamichos D. Corneal injury: Clinical and molecular aspects. Experimental Eye Research 2019;186.

37. Kumari SS, Varadaraj M, Menon AG, Varadaraj K. Aquaporin 5 promotes corneal wound healing. Experimental Eye Research 2018;172:152–158.

38. Ljubimov AV, Saghizadeh M. Progress in corneal wound healing. Progress in Retinal and Eye Research 2015;49:17–45.

39. Fini ME, Stramer BM. How the cornea heals - Cornea-specific repair mechanisms affecting surgical outcomes. Cornea 2005;24:S2–S11.

40. Klenkler B, Sheardown H, Jones L. Growth factors in the tear film: Role in tissue maintenance, wound healing, and ocular pathology. Ocular Surface 2007;5:228–239.

41. Schumann S, Dietrich E, Kruse C, Grisanti S, Ranjbar M. Establishment of a Robust and Simple Corneal Organ Culture Model to Monitor Wound Healing. Journal of Clinical Medicine 2021;10.

42. Castro N, Gillespie SR, Bernstein AM. Ex Vivo Corneal Organ Culture Model for Wound Healing Studies. Jove-Journal of Visualized Experiments 2019.

43. Girolamo N. Moving epithelia: Tracking the fate of mammalian limbal epithelial stem cells. Progress in Retinal and Eye Research 2015;48:203–225.

44. West JD, Dora NJ, Collinson JM. Evaluating alternative stem cell hypotheses for adult corneal epithelial maintenance. World Journal of Stem Cells 2015;7:281–299.

45. Seitz B, Hayashi SJ, Wee WR, LaBree L, McDonnell PJ. In vitro effects of aminoglycosides and fluoroquinolones on keratocytes. Investigative Ophthalmology & Visual Science 1996;37:656–665.

46. Milazzo G, Leonardi A, Fregona I, Violato D, Papa V. Effect of netilmicin and ofloxacin on human keratocytes in vitro. Investigative Ophthalmology & Visual Science 2004;45:U321–U321.

47. Agarwal P, Rupenthal ID. In vitro and ex vivo corneal penetration and absorption models. Drug Delivery and Translational Research 2016;6:634–647.

48. Moreira LB, Lee RF, deOliveira C, LaBree L, McDonnell PJ. Effect of topical fluoroquinolones on corneal re-epithelialization after excimer laser keratectomy. Journal of Cataract and Refractive Surgery 1997;23:845–848.

49. Fukuda M, Inoue A, Sasaki K, Takahashi N. The effect of the corneal epithelium on the intraocular penetration of fluoroquinolone ophthalmic solution. Japanese Journal of Ophthalmology 2004;48:93–96.

50. Carson DL, Chandler M, Hackett R, Edelhauser HF. Ocular toxicity of Ciprofloxacin/PSSA fluoroquinolone antibacterial solution in pigmented rabbits. Journal of Toxicology-Cutaneous and Ocular Toxicology 1996;15:165–178.

51. McDonald EM, Ram FSF, Patel DV, McGhee CNJ. Topical antibiotics for the management of bacterial keratitis: an evidence-based review of high quality randomised controlled trials. British Journal of Ophthalmology 2014;98:1470–1477.

52. Patel GM, Chuang AZ, Kiang E, Ramesh N, Mitra S, Yee RW. Epithelial heating rates with topical ciprofloxacin, ofloxacin, and ofloxacin with artificial tears after photorefractive keratectomy. Journal of Cataract and Refractive Surgery 2000;26:690–694.

53. Wilhelmus KR, Abshire RL. Corneal ciprofloxacin precipitation during bacterial keratitis. American Journal of Ophthalmology 2003;136:1032–1037.

54. Hendrix DVH, Ward DA, Barnhill MA. Effects of antibiotics on morphologic characteristics and migration of canine corneal epithelial cells in tissue culture. American Journal of Veterinary Research 2001;62:1664–1669.

